# Graph Network-Based Analysis of Disease-Gene-Drug Associations: Zero-Shot Disease-Drug Prediction and Analysis Strategies

**DOI:** 10.1101/2024.12.30.630746

**Authors:** Yinbo Liu, Guodong Niu, Siqi Wu, Jingmin Wang, Hesong Qiu, Wen Zhang

## Abstract

Existing drug repurposing methods have key limitations, primarily stemming from their reliance on known direct associations between diseases and drugs for supervised learning, as well as the need for large amounts of prior disease or drug information or feature data. In practice, many disease-drug connections remain unknown, and prior information is often complex and difficult to acquire and organize, limiting the applicability of these models. Furthermore, these models generally lack interpretability, making it difficult for experts to assess the reliability of predictions based solely on standard metrics, which raises doubts about the trustworthiness of their results. To address these challenges, we propose ZS-GNT, an innovative new workflow for zero-shot drug repurposing that leverages a novel and ingenious graph data meta-path linking scheme, which does not require any known disease-drug associations or their prior features. This approach is implemented using the Graph Neural Transformer (GNT) algorithm. The method infers disease-drug relationships indirectly through gene action, utilizing disease-gene associations and gene-drug interactions. It also generates a top drug-top gene linkage map, providing clinicians with a visual tool to assess the plausibility of suggested drugs before advancing to clinical trials. Experimental results show that, under the same linking scheme, the GNT algorithm achieved interaction link prediction accuracies of 95.86%, 99.28%, and 99.54% for three diseases, surpassing four other baseline methods. In a test involving a random selection of 100 diseases for drug discovery, among the top 5 recommended drugs from the candidates identified by ZS-GNT from a pool of 33,251 total drugs, the validation rate reached 47.05%, demonstrating the model’s effectiveness in drug discovery.

## 1. Introduction

the relationship between diseases and genes is a vast and rapidly evolving field of study with significant implications for understanding human health [1]. For example, the International Cancer Genome Consortium (ICGC) conducted a comprehensive pan-cancer analysis, revealing novel genomic alterations across different cancer types and showcasing the complex mutational landscape within cancer genomics, marked by a variety of genetic mutations and unique copy number variations. These findings shed light on the molecular mechanisms driving cancer progression and emphasize their close ties to the field of genetics [2, 3]. Furthermore, the impact of rare genetic variations on gene expression and disease outcomes highlights the importance of considering the full spectrum of genetic diversity in disease research [4]. Additionally, linking drug responses to genetic variations, understanding stratified clinical efficacy and safety, rationalizing differences among drugs within the same therapeutic class, and predicting drug efficacy in patient subgroups have become increasingly important [5]. Genome-wide association studies (GWAS) have also been instrumental in identifying critical therapeutic gene targets and pathogenic genes for drug treatments [6]. These pieces of evidence collectively suggest a close interconnection between diseases, genes, and drugs.

The traditional drug discovery process— a cornerstone of pharmaceutical research— is notorious for its high costs, long timelines, and low success rates, particularly in clinical trials [7]. Studies analyzing the high failure rates of Phase III trials and regulatory submissions have revealed the immense complexity involved in developing marketable drugs [8]. Decades of pharmaceutical innovation have demonstrated the urgent need for a paradigm shift to enhance the efficiency of drug discovery and development [9]. These findings collectively underscore the pressing demand for innovative approaches that can overcome existing inefficiencies and increase the likelihood of success in drug development.

Machine learning (ML) and deep learning (DL) have become transformative tools with the potential to revolutionize the drug discovery process [10, 11]. For example, in HIV research, a Long Short-Term Memory (LSTM) variational autoencoder demonstrated high predictive accuracy and generated promising new drug candidates [12]. Other innovations, such as multitask machine learning models capable of predicting various drug properties, have shown potential in streamlining the drug development pipeline [13]. The exponential growth of transcriptomic data has opened new avenues for exploring compound-protein interactions, particularly in the fields of cellular transcriptomics and RNA biology. Deep generative models can also support gene-level response interpretation of diseases based on genetic features and computational drug screening [14]. Graph Convolutional Networks (GCNs) have gained widespread attention as powerful tools for analyzing complex biological networks in bioinformatics [15]. GCNs have proven invaluable in interpreting these complex transcriptomic landscapes, greatly enhancing our understanding of molecular interactions [16, 17]. Significant progress has been made in disease gene discovery [18] and drug discovery [14, 15] using graph neural networks. However, these drug discovery tools all rely on substantial prior knowledge to construct features and direct disease-drug associations [14, 15].

Based on this, we propose ZS-GNT, an innovative method for drug repurposing that enables zero-shot learning and does not require knowledge. The method constructs a meta-path network that connects various components. It first identifies genes known to be associated with the primary disease of interest [19]. Using these genes, the method selects related secondary diseases that share common genetic links. Next, it identifies secondary genes associated with the secondary diseases, thereby expanding the pool of potentially relevant genes. These expanded gene sets are then used to screen for drugs capable of targeting the corresponding gene loci. The resulting meta-path network highlights the relationships between key diseases, their associated genes, related secondary diseases, and potential drug candidates. This process constructs a focused global network of diseases, with millions of edges, with the primary diseas of interest embedded within. We employ a graph neural network transformer to encode and extract useful latent information from this complex information landscape [20, 21], addressing the challenge of inferring indirect relationships between diseases, genes, and drugs. By integrating functional gene data, ZS-GNT can predict disease-drug associations without known samples and generate a comprehensive interaction network linking drugs, genes, and diseases. This network provides clinicians with a valuable resource, enabling them to assess the feasibility of suggested drugs before advancing to clinical trials.

Our research combines bioinformatics with an in-depth analysis of global gene networks, offering a refined approach to constructing a complex map of disease-gene-drug interactions. This methodology facilitates a deeper exploration of these relationships, accelerating the identification of potential drug targets. Furthermore, our computational models aim to streamline the drug development pipeline, addressing the challenges presented by initial research, clinical trials, and regulatory approval. Ultimately, our work bridges the gap between scientific discovery and clinical application, enhancing the efficiency of drug discovery and accelerating the clinical implementation of therapeutic strategies.

## 2. MATERIALS AND METHODS

### 2.1 Data Collection

An important resource supporting the construction of disease-gene networks is the DisGeNET database [22], which serves as a comprehensive discovery platform. DisGeNET integrates data from expert-curated databases and information extracted via text mining, encompassing a multitude for disease data such as Mendelian diseases, complex diseases, and animal disease models. It assigns scores to gene-disease associations based on the quality of supporting evidence, prioritizing relationships with more robust evidence. As of now, the platform includes over 950,000 gene-disease associations, covering more than 26,133 genes and 30,293 diseases, making it one of the most significant resources in the field.

Complementing DisGeNET is the Drug-Gene Interaction Database (DGIdb) 2.0, a web platform dedicated to mining clinically relevant drug-gene interactions. Since its establishment, DGIdb has been significantly expanded in the updated 2.0 version, now integrating data from 27 different sources, with a particular focus on drug-gene interactions related to clinical trials [23]. Currently, it includes 33,251 drugs or compounds, with a total of 85,460 association records.

First, we obtained disease-gene association data from the DisGeNET database (https://www.disgenet.org/home/). This data includes gene identifiers related to the primary disease under study, which are used to construct the disease-gene associations. Next, we downloaded gene-drug interaction data from the DGIdb database (https://www.dgidb.org/downloads), which contains drug concept IDs linked to the relevant genes, forming the gene-drug interaction dataset. Additionally, we downloaded information on known approved drugs from the ChEMBL database (https://www.ebi.ac.uk/chembl/) to serve as a reference standard for drug selection in subsequent analyses [24]. ChEMBL contains 6,393 known drugs and 1,959 diseases with 36,185 clinical applications.

### 2.2 The global network centered around the focus major disease

As briefly described in the introduction, to construct a disease-gene-drug interaction network centered around a major disease, we first selected a primary “focus disease”, or a primary disease of interest, and identified all genes directly associated with it. Then, for each identified gene, we further identified other diseases related to that gene, which we termed secondary diseases. Next, for these newly mapped secondary diseases, we identified additional genes associated with them, referred to as secondary genes, thus forming a meta-path global network involving the key disease, genes, secondary diseases, secondary genes, and drugs. Through this process, we have built a graph network relationship with millions of edges connecting the “focus disease” to other associated genes and diseases. This network effectively captures both direct and indirect disease-gene association information as we have integrated all relationships involving the focus disease (its related diseases, and associated genes) into a unified data network, ultimately providing a multi-layered perspective that goes beyond direct gene relationships to the focus disease. This integration of disease-gene associations and gene-drug interactions laid the data foundation for our zero-shot and prior knowledge-free predictive analysis. Together, these databases and tools establish a solid foundation for the application of GNT in analyzing disease-gene-drug interactions through streamlining data integration and exploration (**Figure 1**).

**Figure 1.**
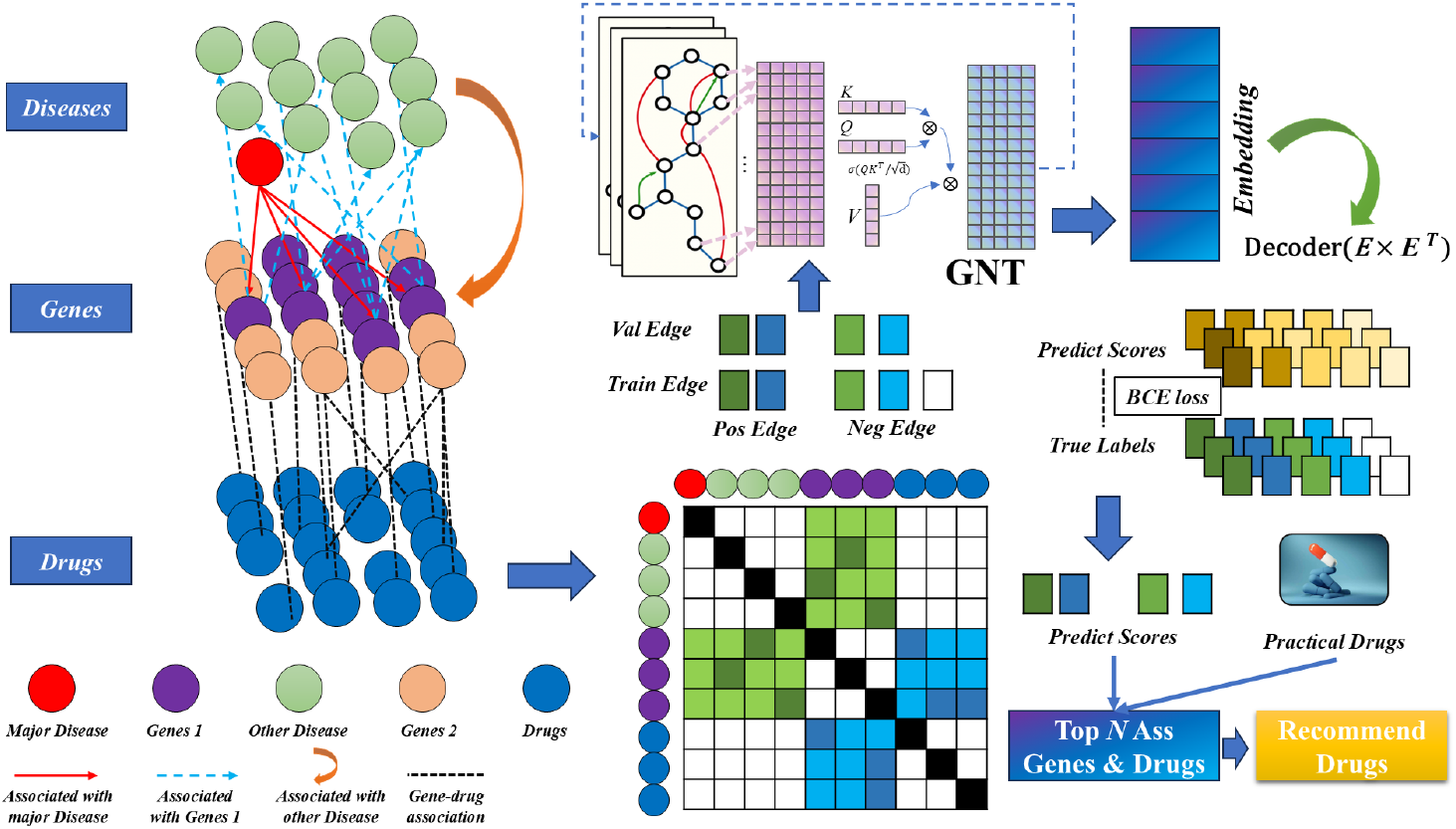
Disease-Gene-Drug association analysis flowchart based on ZS-GNT

Based on the constructed ‘major disease’ (disease, gene, drug) relationship data, we merge them to form a homogeneous graph. The nodes representing diseases, genes, and drugs are denoted as *N*_*D*_*is, N*_*G*_, *E*_*D*_*r*, respectively. Therefore, all nodes in the graph can be represented as:

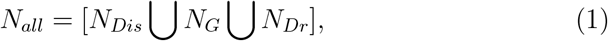

The edges constructed above are denoted as:

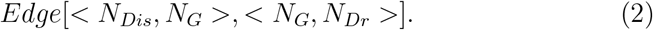

Subsequently, all the edges are integrated into a homogeneous matrix A representing the disease, gene, and drug relationships:

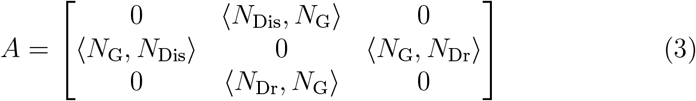

Then, we created an feature matrix for all nodes using one-hot encoding.

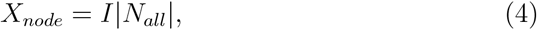

then the comprehensive graph G is denoted as:

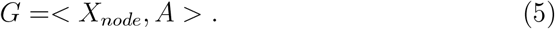

### 2.3 GNT (Graph Neural Network Transformer) Method

The Graph Neural Network Transformer consists of graph convolutional layers, transformer layers, and residual connections. For the graph convolution at the l-th layer, the output is:

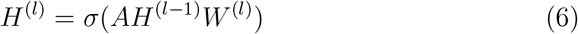

where *W* ^(*l*)^ is the learnable weight matrix at the l-th layer, *σ* is the activation function, and *H*^(1)^ = *X*_*node*_. The input feature matrix *H*^(*l*)^ is used to construct the self-attention mechanism. First, the query (Query), key (Key), and value (Value) are computed as follows

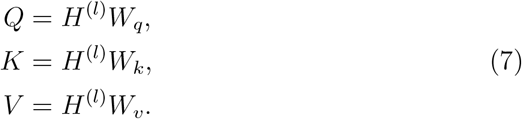

where *W*_*q*_, *W*_*k*_, *W*_*v*_ are the learnable weight matrices. The expression for the Transformer can be represented as:

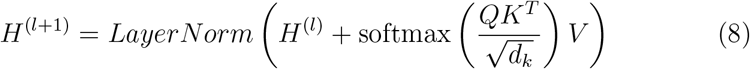

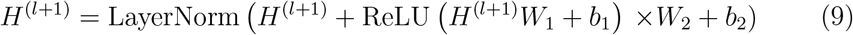

where *d*_*k*_ is the dimension of the key, used for scaling to ensure stable gradients. The output of each layer is accumulated through a residual connection:

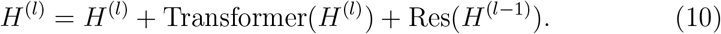

### 2.4 Interaction link prediction

For link prediction, the input features are:

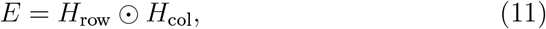

the final output score is:

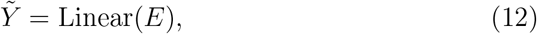

Where ⊙ denotes element-wise multiplication, and 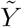 is the predicted link score. This score represents the disease-gene links, gene-disease interaction links, along with their corresponding negative edges, as well as the missing links in the homogeneous graph A. Therefore, 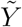 can be used to construct the overall predicted score matrix 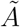 for A.

### 2.5 Loss function

In this model, we adopt the cross-entropy loss function as the primary criterion for evaluating the model’s performance [25].

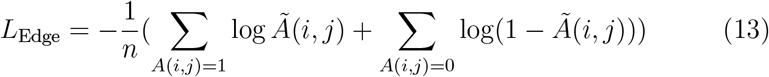

where *L*_Edge_ represents the loss function for the overall loss of the entire homogeneous graph, where n denotes the total number of samples. *A*(*i, j*)= 1 indicates an association between source and target for node i and node j. *A*(*i, j*) = 0 indicates that there is no association between node i and node j in. In addition, we update the parameters of our model using the Adam optimization algorithm [26].

### 2.6 Focus Major Disease Drug Prediction

We select the top genes associated with a specific disease from the predicted disease-gene scores:

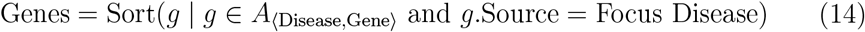

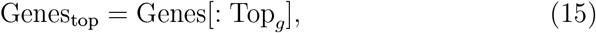

For each top gene, we identify the related top drugs:

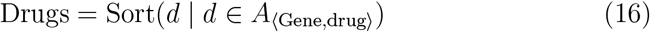

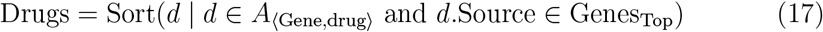

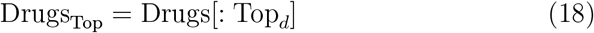

Then, for each Drugs_Top_, we count the corresponding Genes_top_ in the entire filtered data and reorder them based on the count:

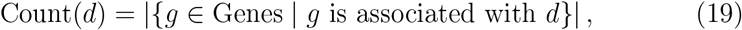

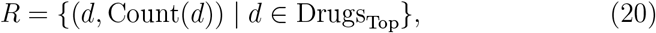

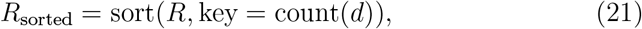

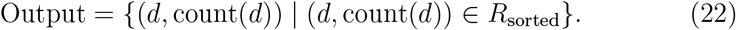

### 2.7 Evaluation Metrics

To comprehensively evaluate the performance of the model, we adopted a series of evaluation metrics, including accuracy, precision, recall, F1 score, and specificity to comprehensively measure model performance based on diverse criteria. These metrics are calculated as follows [27]:

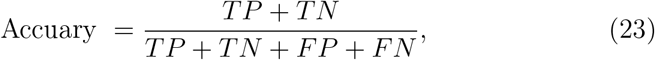

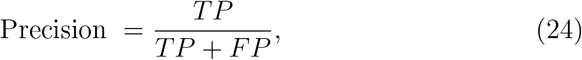

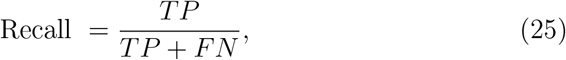

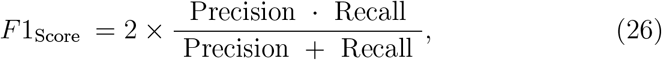

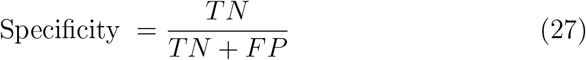

In the above equation, everything is based on the counts of true positives (TP), true negatives (TN), false positives (FP), and false negatives (FN).

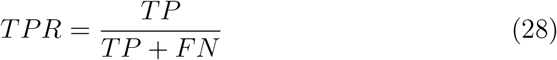

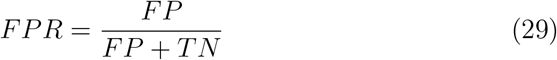

Furthermore, we use the Receiver Operating Characteristic curve (ROC curve), a graphical tool used to evaluate a binary classification model, to further visualize model performance. It plots the False Positive Rate (FPR) on the x-axis and the True Positive Rate (TPR) on the y-axis. The area under the ROC curve is known as AUC (Area Under the Curve), and the closer the AUC value is to 1, the better the model’s performance. We also use the Precision-Recall curve (PR curve) to plot the precision on the x-axis and recall on the y-axis. We denote area under the PR curve as the AUC-PR (henceforth called PRC for conciseness). We will use the AUC and PRC to validate the model’s performance. To assess the matching between the recommended drugs and the actual clinical drugs, to consider the fact that the model may uncover potential drug paths that are completely unknown in clinical settings previously, we introduce the Drug Existence Ratio (*Ratio*_*DE*_) as an evaluation metric. The *Ratio*_*DE*_ is calculated as follows:

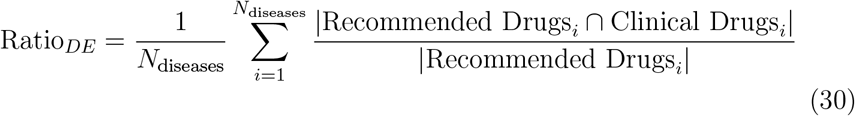

where, *N*_*diseases*_ is the total number of selected diseases, and |·| denotes the number of elements in a set.

## 3. RESULTS

### 3.1 Case Selection

In our research, we have chosen three diseases which, when the method of building their corresponding disease-gene network is applied, result in a range of network complexities, from the least to the most intricate (**Figure 2**). The most straightforward case is C1333977, hepatitis B virus-related hepatocellular carcinoma, a significant form of liver cancer linked to a single viral agent [28]. Next is C0279672, cervical adenocarcinoma, a cancer triggered by viruses such as HPV, which implicates a multitude of genes and has a more extensive gene interaction network [29]. Lastly, C0333806, psoriasis—an immunologically mediated disease—features an extensive gene association network, making it an ideal model for studying genetic interactions in complex diseases. [30].

**Figure 2.**
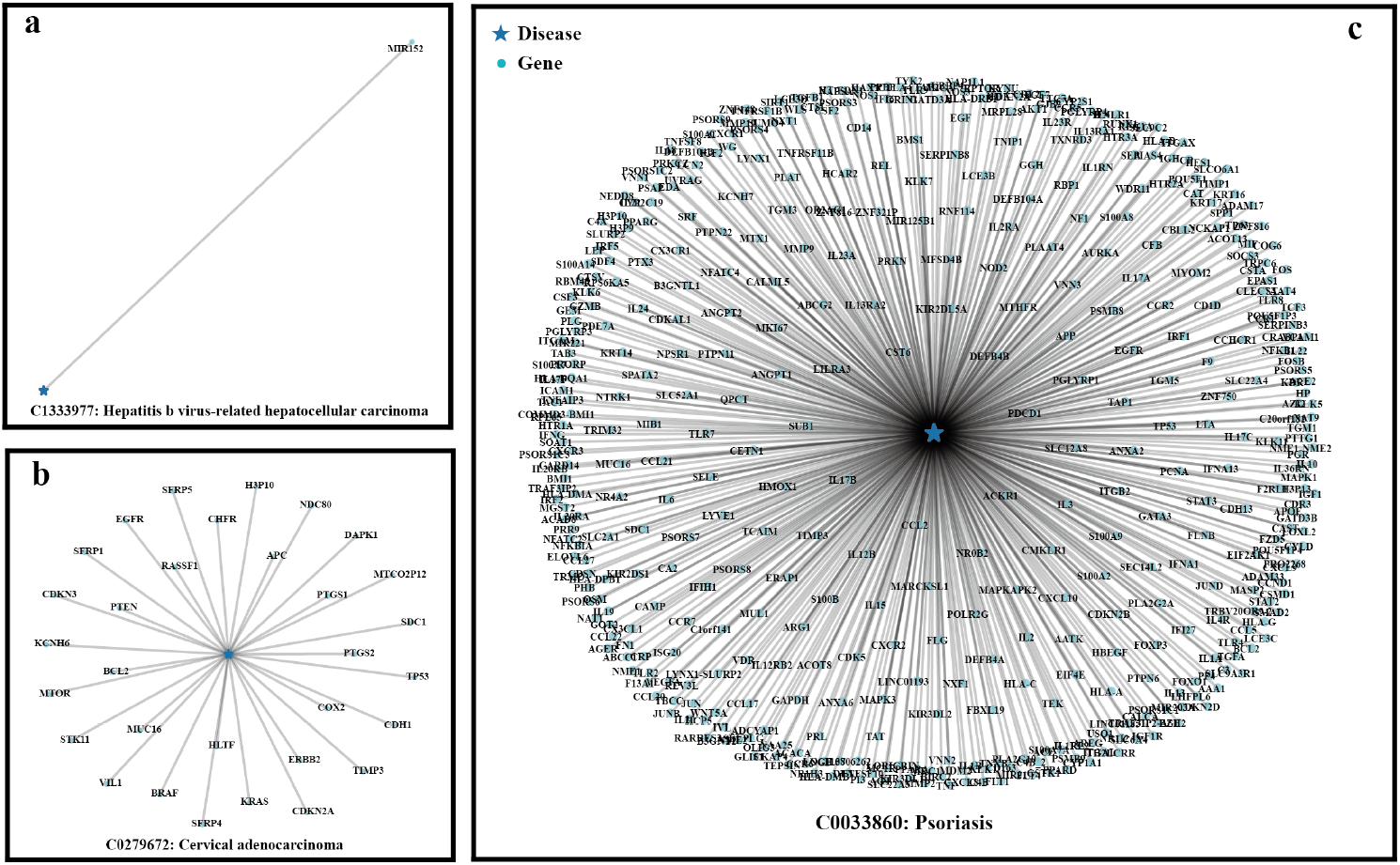
Complexity Analysis of Disease Gene Network Structures and Linkage Relationships (a: C1333977, b: C0279672, c: C0333806)

Visual representations of these networks are depicted above. C1333977 is only associated with a single gene node MM1422, representing the simplest network structure (**Figure 2a**). As for C0279672, the disease node is located centrally, with multiple gene nodes surrounding it, forming a starshaped structure, reflecting a medium-sized disease-gene association network (**Figure 2b**). Lastly, panel C shows the connection between C0333806 and numerous genes, forming a dense network structure, reflecting the complexity of the disease (**Figure 2c**).

**Table 1:** Statistical Table of Disease-Related Data Information.

| CID | Disease Name | N<sup>1</sup> | N<sup>2</sup> | N<sup>3</sup> | N<sup>4</sup> | N<sup>5</sup> |
| --- | --- | --- | --- | --- | --- | --- |
| C1333977 | HBV<sup>a</sup> | 1 | 13 | 6872 | 9638 | 187143 |
| C0279672 | CAC<sup>b</sup> | 31 | 3050 | 12984 | 10679 | 5287751 |
| C0333806 | Psoriasis | 471 | 7354 | 13640 | 10691 | 6331044 |

1. Number of Associated Primary Genes; 2. Number of Diseases Associated with Primary Genes; 3. Number of Associated Secondary Genes; 4. Number of Associated Drugs; 5. Total Number of Edges; - a. Hepatitis B virus-related hepatocellular carcinoma; - b. Cervical adenocarcinoma.

### 3.2 Drug Analysis for Disease Treatment Based on Model Predictions

The figure shows the top 10 drugs predicted by the model for diseases such as (a: C1333977, b: C0279672, c: C0333806), highlighting potentially effective candidates for treatment (**Figure 3**).The numbers under the drug nodes represent the number of associated key genes. These results provide a new perspective for optimizing treatment plans for diseases and can help clinicians make more precise drug selections based on the model’s predicted list of drugs. Analysis of the predicted drugs reveals that they comprise of anti-inflammatory drugs, immunosuppressants, or anticancer drugs, which are closely related to the pathological mechanisms of the focus diseases, indicating the correctness of the results. Furthermore, these predictions reveal that some drugs may be more effective in treating specific diseases in cases where the treatment outcomes have not been fully validated in the real world. This serves as valuable leads for future drug research and clinical trials. Moreover, we have provided gene-linkage visualizations for the drugs, which increase the interpretability of drug effects. This process helps doctors make better-informed drug choices based on their clinical experience.

**Figure 3.**
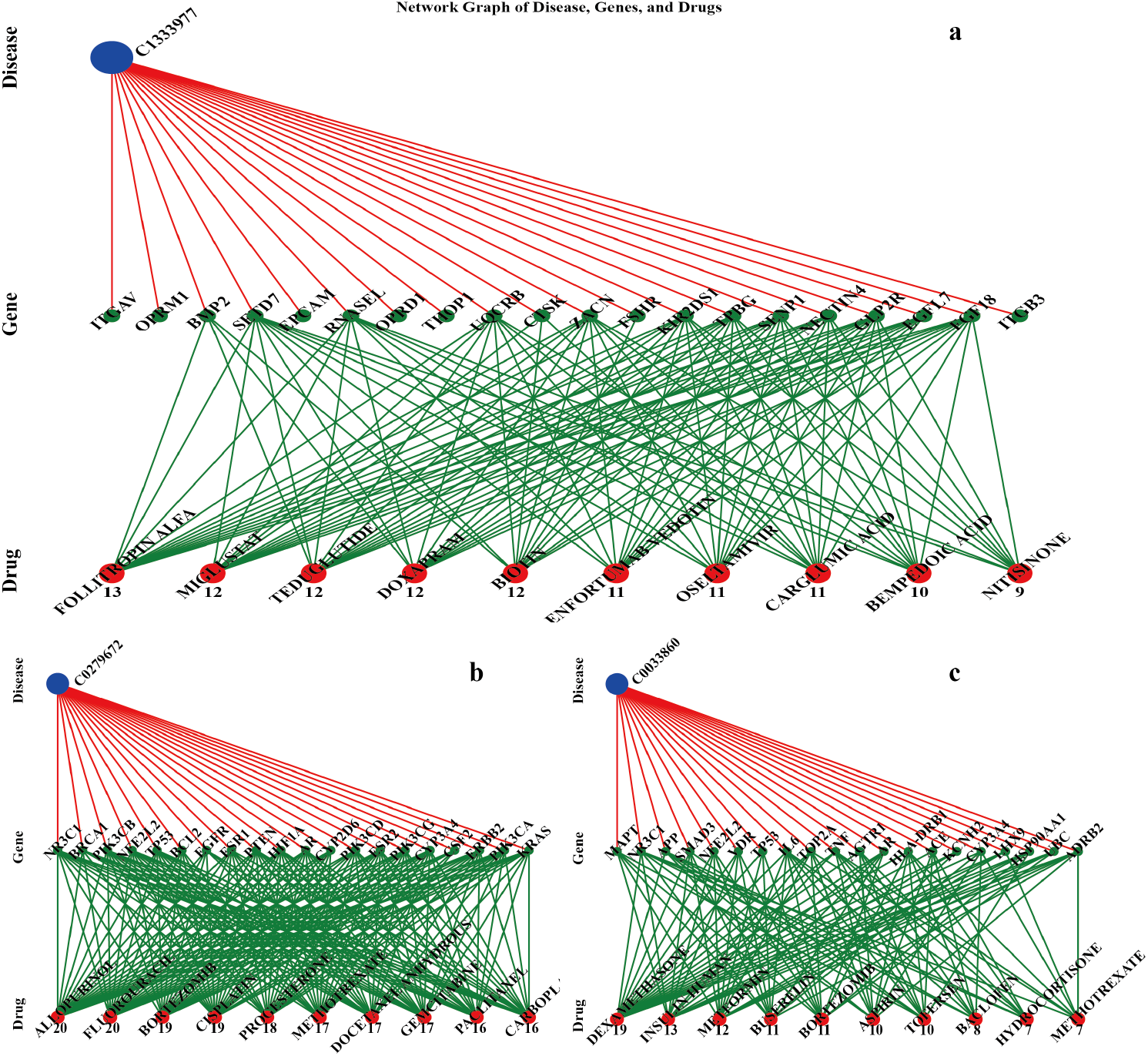
Top 10 drugs for treating diseases predicted by the model(a: C1333977, b: C0279672, c: C0333806).

We provide examples of real-world implications of our work below. In the results for C1333977 presented in Figure 3, it was found that although ITGAV is the main key gene [31], it is not a very good drug target. Slightly lower ranked genes such as BMP2 [32] and SETD7 [33], however, are genes that can be targeted by drugs and may be more relevant to drug discovery.

The drug that targets the most key genes, Follitropin alfa, is a recombinant human follicle-stimulating hormone (rFSH) primarily used to promote ovarian function in women and is commonly used in the treatment of infertility [34]. In a 2022 medical record, ovarian stimulation was noted to increase viral replication of HBV [35]. This new discovery suggests that this drug may be useful for treating C1333977 symptoms. Experts can assess the reliability of this prediction based on the target genes.

Similarly, for the diseases C0279672 and C0333806, the corresponding key genes and their associated drugs were identified in Figure 3, with the top recommended drugs being Allopurinol and Dexamethasone, respectively. Allopurinol is a drug used to treat hyperuricemia, primarily by inhibiting xanthine oxidase, a key enzyme in purine metabolism, to lower uric acid levels in the body [36]. It has been experimentally combined for treating ACA [37, 38]. Dexamethasone is a corticosteroid drug with strong antiinflammatory, immunosuppressive, and anti-allergic effects [39]. It works by binding to steroid receptors in the body to suppress inflammation and regulate the immune system. Numerous studies have shown that this drug can be highly effective in treating psoriasis [40, 41], further validating the model results.

### 3.3 Comparison with Other Methods

To test the performance of the GNT network, we selected four graph neural network methods (GIN [42], GAT [43], GCN [44], GraphSAGE [45]) as baselines and illustrated the performance of GNT and these methods across different metrics (Precision, Recall, Specificity, AUC, PRC) using 3D bar plots (see **Figure 4**). On the three different drug-gene datasets (C1333977, C0279672, C0033860) mentioned above, GNT demonstrates significant performance gains across multiple metrics. Specifically, GNT achieved AUC values of 95.86%, 99.28%, and 99.54%, and AUPRC values of 95.56%, 99.24%, and 99.50% (refer to **Supplementary Tables 1-3**), validating the model’s ability to dicipher networks at a global scale and encode their features. Methods such as GIN and GAT perform well in specific metrics (e.g., recall or specificity), but their overall performance may not be as reliable as GNT. The strong performance of GNT on AUC and PRC, which are comprehensive evaluation metrics, highlights the superiority of our method in a broader context. Furthermore, a comparison of the subfigures (a, b, c) shows that the GNT model consistently performs well across all datasets, while baseline methods exhibit greater variability, indicating that our method is more generalizable.

**Figure 4.**
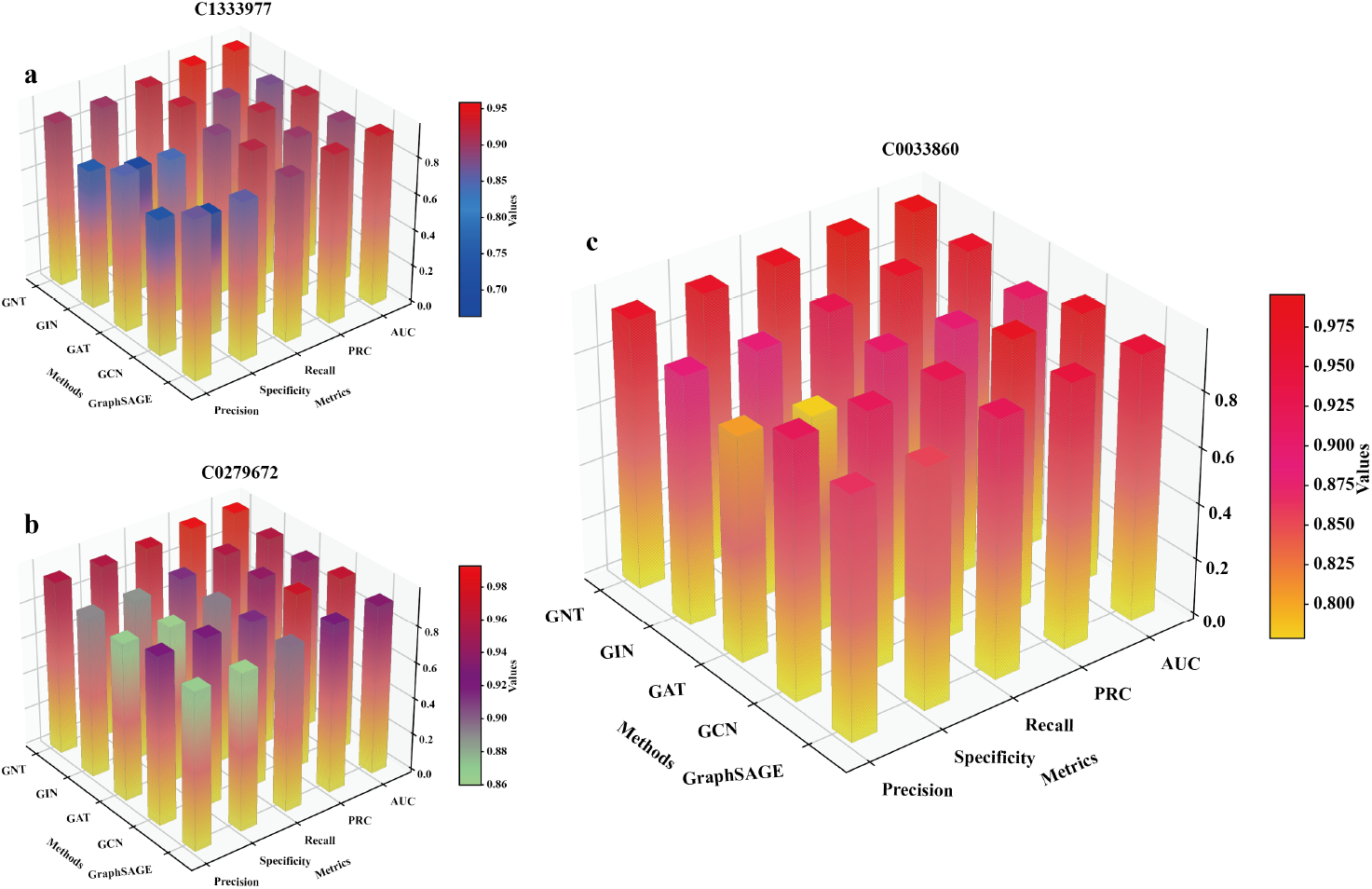
Comparison of network performance between GNT and other methods

### 3.4 Model Analysis

#### 3.4.1 Analysis of the impact of model architecture and regularization

The choice of hyperparameters plays a critical role in the training process of machine learning models. Therefore, we provide a thorough and systematic analysis of key hyperparameters: the number of hidden channels, the number of GCN layers, and the dropout rate, and quantify their effect on model performance (**Figure 5**).

**Figure 5.**
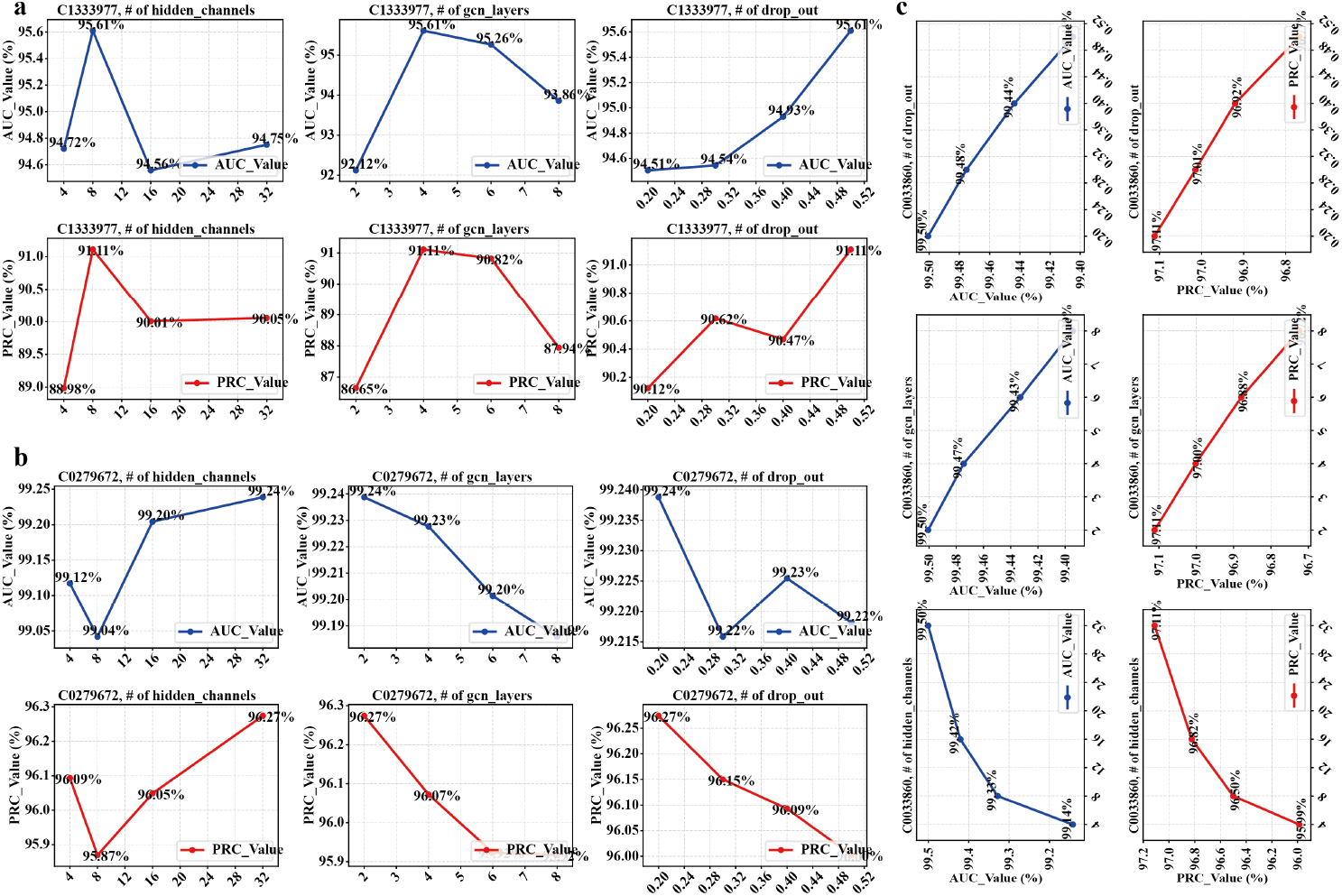
Hyperparameter Sensitivity Analysis Chart

In the experiments with models trained for C1333977 and C0279672, increasing the number of hidden channels significantly improved the AUC, indicating that more feature channels helped the model extract richer feature information. However, as the number of hidden channels increased further, the AUC decreased, and the PRC value also displayed a downward trend. These results suggest that overfitting may have occurred as the model’s complexity grew, degrading the precision and recall.

Increasing the number of GCN layers generally improves AUC, indicating better understanding of the graph’s structure. However, PRC decreases with more layers, suggesting overfitting due to model complexity. Moderate dropout rates boost AUC, especially for C1333977 and C0279672 models, enhancing generalization. However, PRC declines with higher dropout rates, particularly beyond a certain threshold, as excessive dropout may reduce model expressiveness. In summary, while more layers and dropout help improve AUC, they negatively affect PRC when overdone. Hyperparameter optimization should balance AUC with precision and recall.

#### 3.4.2 Analysis of the impact of activation functions

This figure compares model performance using different activation functions, specifically analyzing the ROC and PRC curves. The three subplots above demonstrate the results of different activation functions on three datasets: C1333977, C0279672, and C0333806 (**Figure 6**).

**Figure 6.**
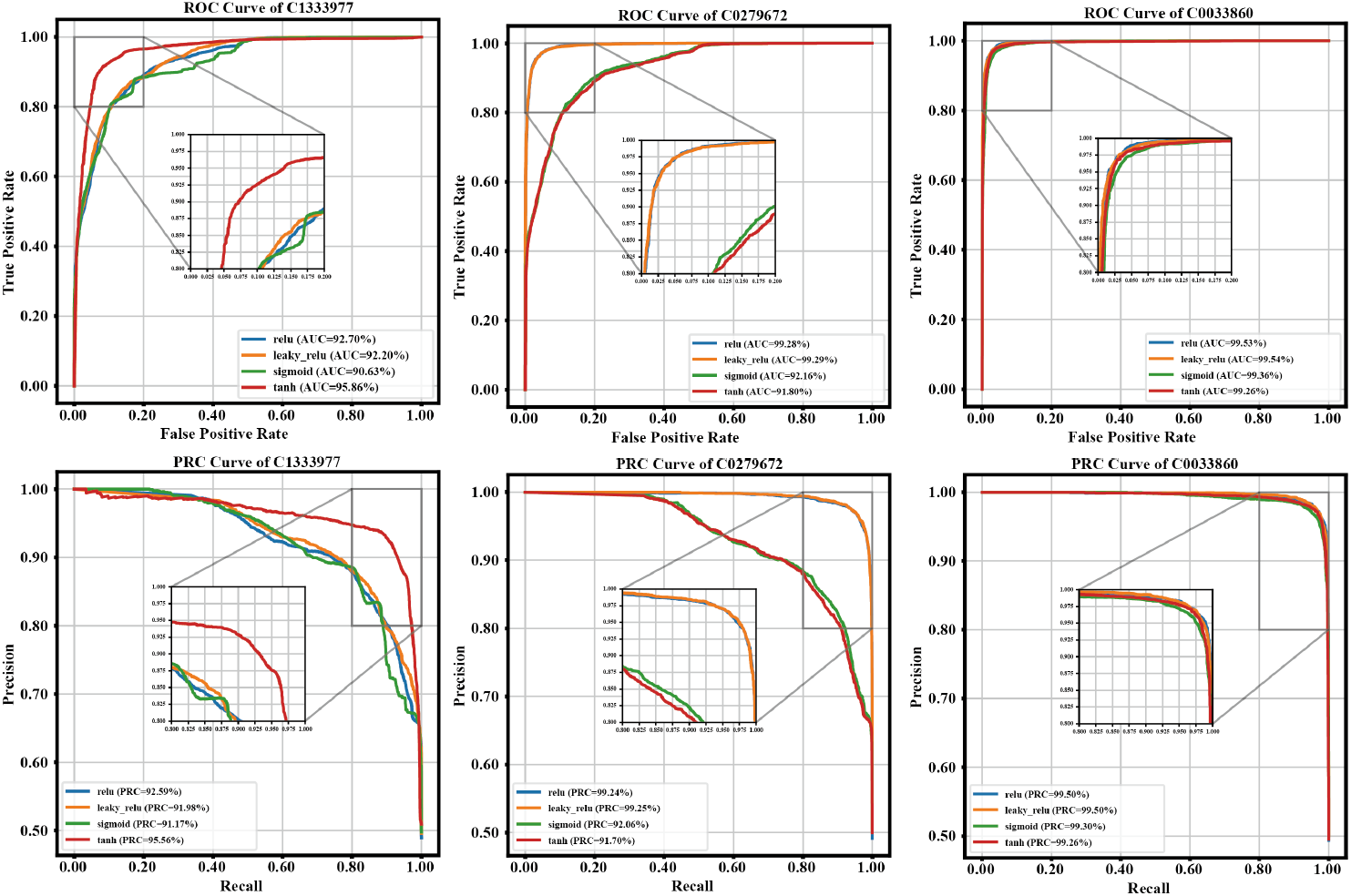
Analysis of the Impact of Activation Functions on Model Performance – ROC and PRC Curve Graphs

Analyzing the C1333977 dataset, we observe the performance difference between four activation functions (ReLU, leaky ReLU, sigmoid, tanh) on the ROC and PRC curves. The ReLU activation function outperforms the others, achieving the highest AUC and PRC values of 99.79% and 99.53%, respectively. In contrast, despite similar PRC values, the sigmoid and tanh functions perform poorly, with their AUC values being significantly lower. Leaky ReLU performs well but ultimately falls short of ReLU. ReLU consistently outperforms the other activation functions across all datasets. For the C0279672 dataset, ReLU achieves the highest AUC (99.29%) and PRC (99.55%), slightly surpassing leaky ReLU and tanh. Sigmoid has the lowest AUC (99.15%), but its PRC is similar to the others. On the C0333806 dataset, ReLU leads with an AUC of 99.28% and PRC of 99.25%, followed closely by leaky ReLU. Sigmoid and tanh perform poorly, especially in the PRC curve. In summary, ReLU is the most optimal activation function for AUC and PRC, with leaky ReLU also performing well. Sigmoid and tanh show weaker performance overall.

#### 3.4.3 Analysis of the impact of training epochs

Training time is a critical factor in model learning. Hence, we further study the trend of AUC and PRC values across training epochs for different models (C1333977, C0279672, C0333806) (**Supplementary data**).

During training, AUC values initially improve but gradually slow down and stabilize, suggesting that the model may have reached its performance limit, possibly due to factors such as learning rate decay or overfitting. PRC values also increase in the early stages, indicating a good balance between precision and recall, but may decline later on. Overall, while AUC improves early on, it gradually plateaus, and PRC improves initially before declining, highlighting the need for careful management during training to prevent overfitting and ensure a balance between AUC and PRC.

### 3.5 Validation of ZS-GNT Results on Clinically Approved Drugs

To validate the potential clinical applications of the ZS-GNT model, as shown in Figure 7(a) we randomly selected 100 diseases from a total of 30,293 diseases or conditions in the DisGeNET database for investigation. Among these 100 selected diseases, 63 had known gene associations, meaning that they could be used by ZS-GNT for disease recognition by applying the methods in the paper. Out of these 63 diseases, 17 were present in the ChEMBL database.

**Figure 7.**
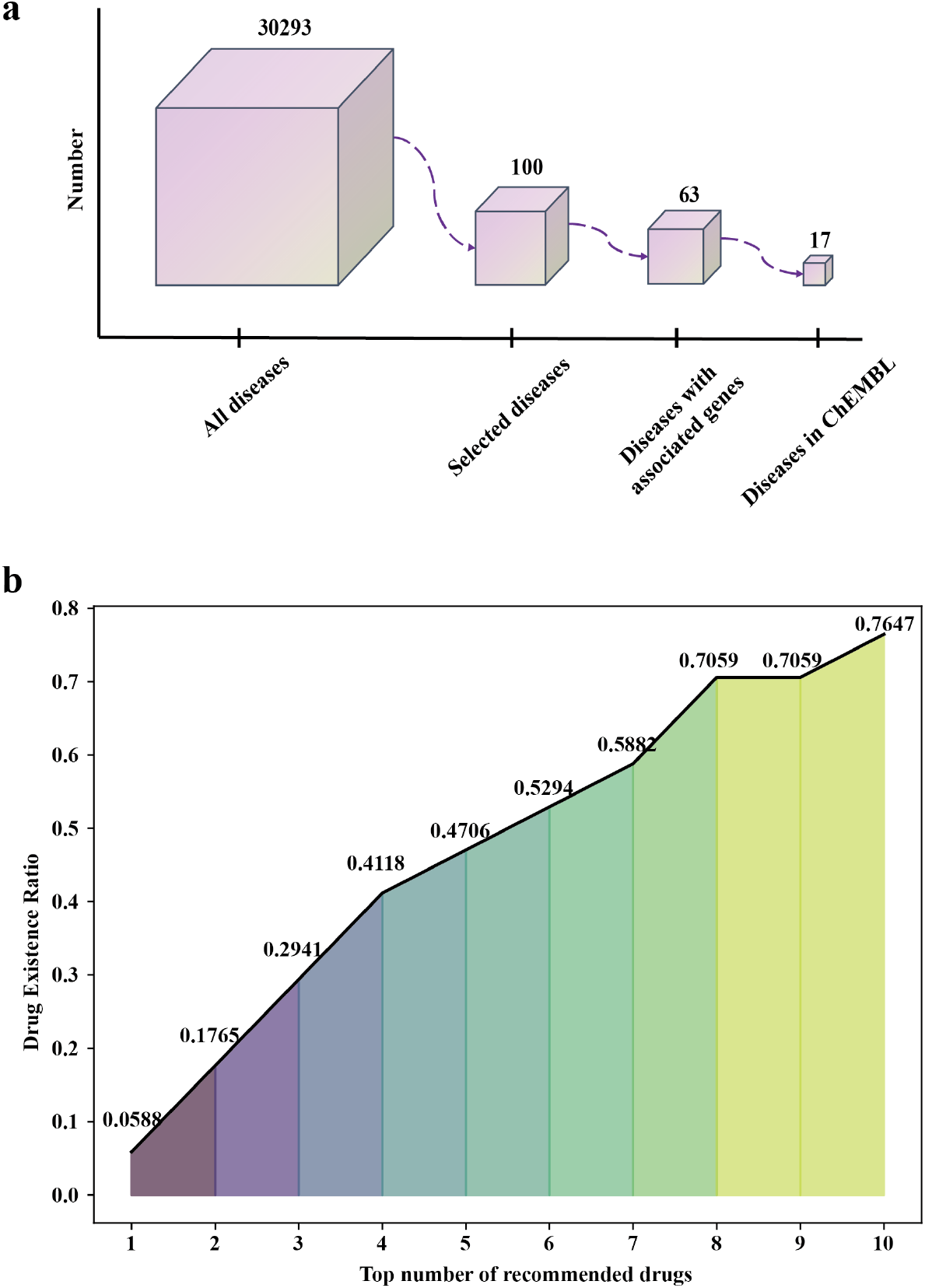
Analysis of the Impact of Activation Functions on Model Performance – ROC and PRC Curve Graphs

For drug recommendation, we used the DGIdb database and applied ZS-GNT to predict the top 10 recommended drugs for each of the 63 diseases, along with their potential target genes. Detailed information can be found in the appendix. Moreover, we calculated the Drug Existence Ratio (equation (29)) for the recommended drugs of the 17 diseases that are listed in the ChEMBL database, hence providing a comparision to clinically approved drugs. As shown in Figure 7(b), the Drug Existence Ratio for the top 5 recommended drugs was 47.06%. This means that, on average, there is a 47.06% chance that the top 5 recommended drugs for the diseases in the study are already approved for clinical use. For the top 10 recommended drugs, this ratio increases to 76.47%, indicating that a significant portion of the recommended drugs are already in clinical practice. These results strongly support the clinical validity of the ZS-GNT model.

## 4 DISCUSSION

In this study, we introduced ZS-GNT, a zero-shot analytical approach based on Graph Convolutional Transformers, designed to uncover complex relationships among diseases, genes, and drugs. Our findings provide new insights into zero-shot drug discovery, disease mechanisms, and drug repurposing. By constructing a multi-level graph structure, we developed a comprehensive network centered on major diseases, containing millions of edges and a wealth of latent information. This approach addresses the challenge of relying on large-scale knowledge graph data for constructing prior information, enabling more efficient analysis

The ZS-GNT model demonstrated its ability to identify key genes crucial for the onset, progression, and development of specific diseases through zero-shot disease-drug data analysis. Leveraging GNTs’ inherent capability to analyze graph structures, the study captured gene interactions and regulatory relationships, enabling precise selection of genes with significant roles in disease networks. For instance, in the analysis of neurodegenerative diseases, the model identified genes associated with neuroinflammation, whose abnormal expression has been shown to drive disease progression. This showcases the model’s ability to prioritize biologically relevant genes with clinical significance.

Our GNT model demonstrated strong potential for identifying drugs to treat specific diseases by extracting gene-drug relationships embedded within the graph structure. By targeting key genes and their associated signaling pathways, the predicted drugs showed promise in effectively modulating the expression of disease-related genes. These findings highlight new opportunities for drug repurposing, where existing drugs can be adapted for new therapeutic applications. This approach has the potential to significantly reduce drug development costs and timelines, accelerating the path to clinical implementation.

Cross-validation and independent test set evaluations confirmed the ZS-GNT model’s high accuracy in analyzing disease-gene-drug association networks. Compared to other graph neural network methods, such as GCN and GAT, the ZS-GNT model outperformed in multiple metrics. These results underscore the advantages of ZS-GNT in analyzing complex, structured, and heterogeneous biological networks, further establishing its practical application potential in bioinformatics.

One of the notable outcomes of this study was the identification of previously underexplored or undiscovered disease-drug pairs. Although these pairs are rarely reported in the current medical literature, the model predicted a therapeutic connection between them with strong supporting evidence. For example, in certain rare diseases, the study indicated that abnormalities in specific genes could potentially be mitigated by regulating targeted drugs, providing new insights and hope for treatment. Furthermore, by comparing the predictions with real clinical drug data, the effectiveness of zero-shot predictions for identifying disease-treatment drugs was validated.

Overall, this study demonstrates the effectiveness of our model in discovering and predicting disease-drug relationships. These findings may offer valuable perspectives for understanding the molecular mechanisms of diseases, supporting new drug development efforts, and advancing precision medicine. Looking ahead, as more high-quality biological data becomes available and model algorithms continue to be optimized, ZS-GNT may play a pivotal role in biomedical research. This progress will drive advancements in disease diagnosis, treatment strategies, and the discovery of innovative therapies, ultimately improving patient outcomes.

## ACKNOWLEDGMENTS

This project was supported by the Genoimage R&D:20240703. And we express our deepest gratitude for the support from JIYING TECHNOLOGY CO., LIMITED.

## Code Availability

The source code for our proposed method, ZS-GNT, is publicly available at: https://github.com/yinboliu-git/ZS-GNT. The repository contains the full implementation of our approach, including dataset preprocessing, model training, and evaluation scripts. Detailed instructions for installation and usage are provided in the README file.

